# Getting there in one piece: The Rac pathway prevents cell fragmentation in a nonprotrusively migrating leader cell during organogenesis

**DOI:** 10.1101/2023.12.01.569642

**Authors:** Noor Singh, Karen Jian Li, Kacy Lynn Gordon

## Abstract

The *C. elegans* hermaphrodite distal tip cell (DTC) leads gonadogenesis. Loss-of-function mutations in a *C. elegans* ortholog of the Rac1 GTPase (*ced-10*) and its GEF complex (*ced-5*/DOCK180, *ced-2*/CrkII, *ced-12*/ELMO) cause gonad migration defects related to directional sensing; we discovered an additional defect class of gonad bifurcation in these mutants. Using genetic approaches, tissue-specific and whole-body RNAi, and *in vivo* imaging of endogenously tagged proteins and marked cells, we find that loss of Rac1 or its regulators causes the DTC to fragment as it migrates. Both products of fragmentation—the now-smaller DTC and the membranous patch of cellular material—localize important stem cell niche signaling (LAG-2/DSL ligand) and migration (INA-1/integrin subunit alpha) factors to their membranes, but only one retains the DTC nucleus and therefore the ability to maintain gene expression over time. The enucleate patch can lead a bifurcating branch off the gonad arm that grows through germ cell proliferation. Germ cells in this branch differentiate as the patch loses LAG-2 expression. While the nucleus is surprisingly dispensable for aspects of leader cell function, it is required for stem cell niche activity long-term. Prior work found that *Rac1^−/−^;Rac2^−/−^*mouse erythrocytes fragment; in this context, our new findings support the conclusion that maintaining a cohesive but deformable cell is a conserved function of this important cytoskeletal regulator.

## INTRODUCTION

The formation of each of the two arms of the *C. elegans* hermaphrodite gonad is led by a post-embryonic migration of a somatic gonad leader cell, the distal tip cell (DTC)^1–3^. DTCs begin migrating–nonprotrusively^4^–in opposite directions from the site of the future vulva along the ventral midline at the L2 larval stage. In the L3 larval stage, each DTC executes two 90-degree turns that establish the gonad bend. DTC migration continues along the dorsal body wall during the L3 and L4 larval stages until the two DTCs both reach the dorsal midbody, where they remain throughout adulthood^2,5,6^. Proliferating germ cells provide a propulsive force to the DTC, which interacts with the extracellular matrix to steer the direction of migration^4^.

The distal-most end of the adult germline—capped by the DTC—is composed of germ stem cells followed proximally by their still-mitotic descendants, together comprising the proliferative zone^7^. The DTC acts as the germ stem cell niche by expressing the ligand LAG-2^8^, which is transduced by the GLP-1/Notch receptor in germ cells. Active Notch signaling in the germline leads to post-transcriptional repression of meiotic entry factors to prevent differentiation^9^. Proximal to the proliferative zone, germ cells lose active Notch signaling, lose the repression of meiotic entry factors, enter meiosis, and eventually differentiate as gametes^10–12^. Hermaphrodites make sperm cells in the L4 larval stage and oocytes as adults^13,14^. An essential feature of the adult hermaphrodite germline is that some meiotic cells do not form gametes, but instead take on a nurse-cell-like role in which they donate their cytoplasmic contents to oocytes and undergo apoptosis, becoming engulfed by the somatic gonadal sheath cells that wrap around the germline^15,16^.

Apoptosis and engulfment of dead cells are regulated by highly conserved factors, first identified in a *C. elegans* genetic screen for animals with abnormal cell death (*ced* mutants)^17–25^. Engulfing cells activate two partially redundant *ced* pathways^26^, one of which has also been shown to also be required for DTC migration^20–23^. This pathway is composed of CED-10/Rac1^22^–a regulator of the cytoskeleton^27^ conserved among eukaryotes^28^–and factors that have since been discovered to comprise its upstream activating guanine nucleotide exchange factor (GEF) complex: CED-5/DOCK180^29–31^, CED-2/CrkII^22^, CED-12/ELMO^20,21,30,31^. (CED-12 has also recently been discovered to have GAP function in the context of F-actin nucleation in *C. elegans* embryonic morphogenesis^32^). Rac GTPases, members of the Rho family of GTPases, are required for cytoskeletal remodeling in protrusions at the leading edge of many migratory cells^33,34^. However, the migratory DTC has a smooth leading edge^4,35,36^, both distinguishing it from the classic model of leader cell migration^37–39^ and raising the question of how the Rac-pathway *ced* module–a canonical regulator of lamellipodia–regulates migration in the non-protrusive cell leading gonadogenesis. The DTC may serve as a useful model for the molecular regulation of tubular organs that form with migration of non-protrusive leader cells. One way these Rac pathway *ced* genes regulate DTC migration is as downstream effectors of noncanonical Wnt signaling^40,41^, culminating in repression of the netrin receptor UNC-5 during DTC dorsal migration^40^.

Genetic control of gonadogenesis is well-studied^1,42,43^, and many mutants with polarity defects^40,41,44^ and other gonad malformations have been identified^22,45–51^. Genetic studies of the three *C. elegans* Rac-family GTPase-encoding genes *ced-10*^22^*, rac-2*^52^, and *mig-2*^53^ have revealed that two play a role in DTC migration^22,52,54^. *mig-2*^47,54^ and *ced-10*^22^ cause DTC migration defects when mutated. *ced-10* and *rac-2* are nearly identical at the sequence level^52^ and act in different subsets of cell types (*rac-2* is not expressed in the DTC^55^). How DTC-migration genes act at a cellular level to coordinate the cell movements of organogenesis is an area of increasing interest. Recent work using live-imaging techniques reveals the key role of interactions between the DTC and extracellular matrix, mediated, for example, by integrin^4^ (which in *C. elegans* is composed of INA-1/PAT-3 and PAT-2/PAT-3 dimers^56–58^). We set out to investigate a previously unreported class of gonad defect—bifurcation—in genetic mutants that have long been known to have DTC migration defects.

Here we report that loss-of-function mutations in the Rac-pathway *ced* genes lead to the fragmentation of the DTC as it migrates. Both products of fragmentation—the distal tip cell (reduced in size) and the membranous patch of cellular material—localize important migration (an integrin alpha subunit, INA-1) and signaling (the Notch ligand LAG-2) factors to their plasma membrane. However, only one structure retains the DTC nucleus and therefore the ability to maintain gene expression and synthesize new protein throughout adulthood. The patch can promote local germ cell proliferation for a time, leading to a bifurcating branch off the gonad arm that grows through germ cell proliferation until the patch loses LAG-2 signal due to its lack of nucleus. Cell fragmentation and changes in deformability have been documented in mammalian cell types after Rac1 loss-of-functiony^59–61^; we conclude that Rac1 is required to keep the DTC in one piece during migration, and that the Rac pathway plays a fundamental and broadly-conserved function in maintaining cellular integrity in a number of Eukaryotic cell types.

## RESULTS

### Loss of function of Rac-pathway *ced* genes causes gonad bifurcation

Mutations of the cell death pathway genes *ced-10/*Rac1, *ced-5/*DOCK180, *ced-2/*CrkII, and *ced-12/*ELMO have long been observed to cause gonad migration defects^20–22,29,52^. We examined *ced-10(n1993)*^22^*, ced-5(n1812)*^29^*, ced-2(n1994)*^22^, and *ced-12(n3261)*^21^ mutants, all of which are putative null with the exception of *ced-10(n1993),* a hypomorph^22,52^.

Typically, each gonad arm forms a single U-shaped, blind tube^1,6^. In all four mutants, we observed gonad bifurcation (Figure 1) which was not identified in the original or subsequent reports of gonad defects caused by mutations in *ced* genes (Gumienny et al., 2001, wrote of a different *ced-12(oz167)* allele: “*ced-12* mutants also occasionally show …short branching or distensions”). At the L4 (Figure 1B) and young adult (Figure 1C) stages, we consistently observe bifurcation in 10-16% of animals (Figure 1D). While misdirected DTC migration or turning defects have been investigated in mutants for these^40^ and other genes^35^, gonad bifurcation is strikingly rare^62–64^. We decided to focus on this most severe phenotype to elucidate additional roles of Rac-pathway *ced* genes in regulating DTC migration.

**Figure 1.**
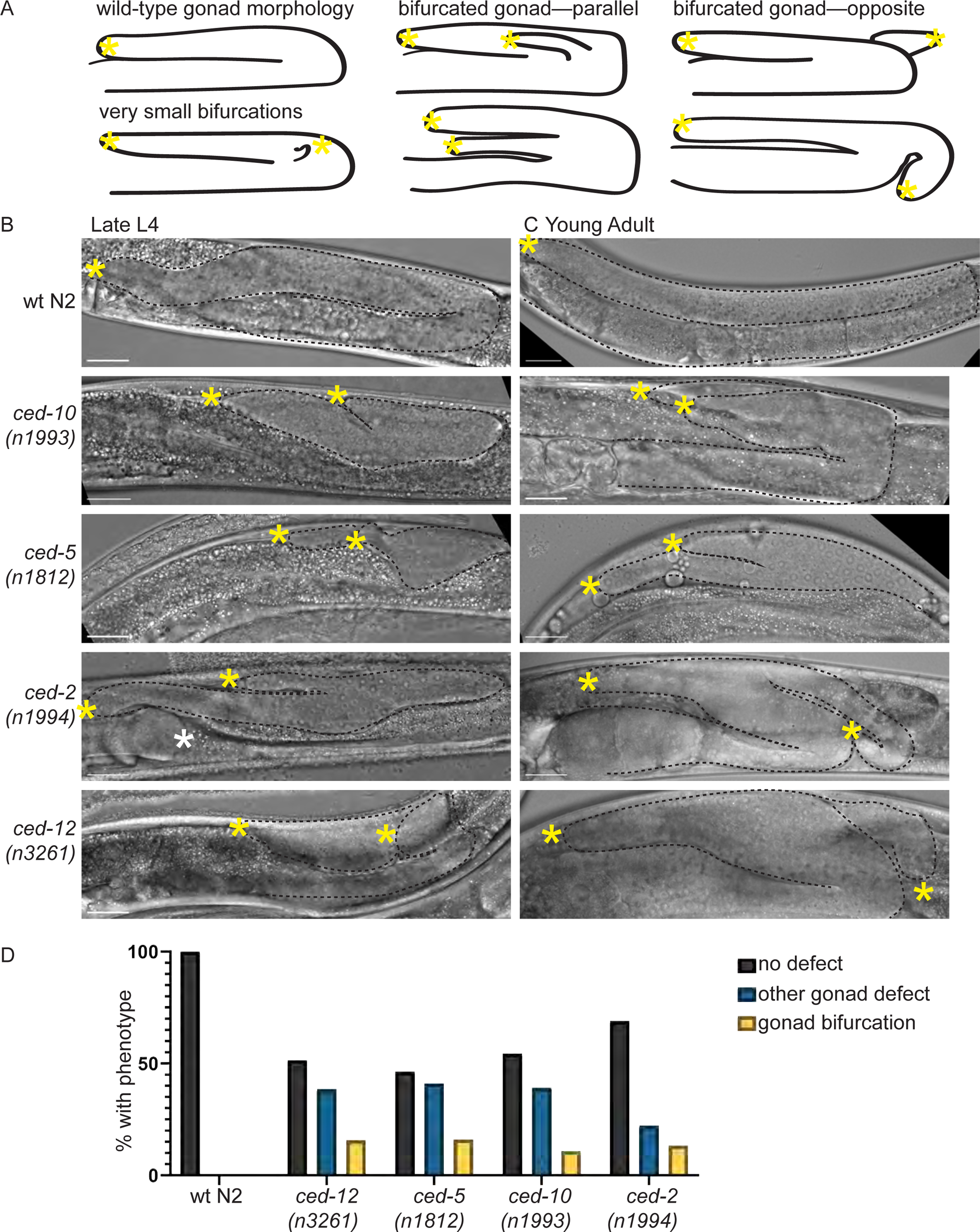
Rac pathway *ced* mutants have gonad bifurcation with moderate penetrance. **(A)** Cartoon illustrating wild-type and bifurcated gonads in *C. elegans* hermaphrodites. (**B)** Micrographs made with DIC imaging of *C. elegans* hermaphrodites in late larval L4 stage for wild-type N2 and four mutant strains. Images are Z-projections through thickness of the gonad required to capture both tips. Visible gonad outlined in black dashed line. Yellow asterisks show the anatomical tips of gonad branches of the gonad on the right-hand side of the image. White asterisk, visible tip of other gonad arm. (**C)** Same as B, but in the young adult stage. Scale bar 20 μm. **(D)** Graph showing percentage of N2 controls and mutants of each genotype (L4s and young adults) with gonad bifurcation (yellow), other gonad defects (blue), or no defect (black). N2 WT control N=51, *ced-5(n1812)* N=56, *ced-10(n1993)* N=46, *ced-12(n3261)* N=70, *ced-2(n1994)* N=45.

### The Rac pathway acts cell-autonomously to regulate DTC leader cell activity, and loss of function causes somatic gonad cell defects as well as gonad anatomy defects

To determine whether gonad bifurcation is caused by cell-autonomous loss of the Rac pathway *ced* genes, we used a strain with DTC-specific sensitivity to RNAi and a distal tip cell membrane marker^65^ (see Methods and Figure 2A) to knock down *ced-10, ced-5, ced-2,* and *ced-12* in the DTC. Indeed, *ced* gene knockdown in the DTC causes cell-autonomous defects that result in bifurcation (Figure 2B). We next examined the DTC itself in RNAi-treated animals. We made two striking observations. First, when the gonad bifurcates, both tips have a focus of mNeonGreen (mNG) signal (Figure 2B), in contrast to the single focus of mNG signal in the control RNAi-treated animals (Figure 2A). Second, even when the gonad does not bifurcate, there is a second patch of mNG signal near the bend of the gonad in 17-29% of L4 larval animals (Figure 2B’). Similar defects were also observed after RNAi of *mig-2*/RhoG, another small Rac-family GTPase, but were not observed after knockdown of the parallel engulfment pathway members *ced-1* and *ced-6* (Figure S1), in agreement with previously observed specificity of the Rac pathway *ced* genes, but not other *ced* genes, for DTC migration^29,52^.

**Figure 2.**
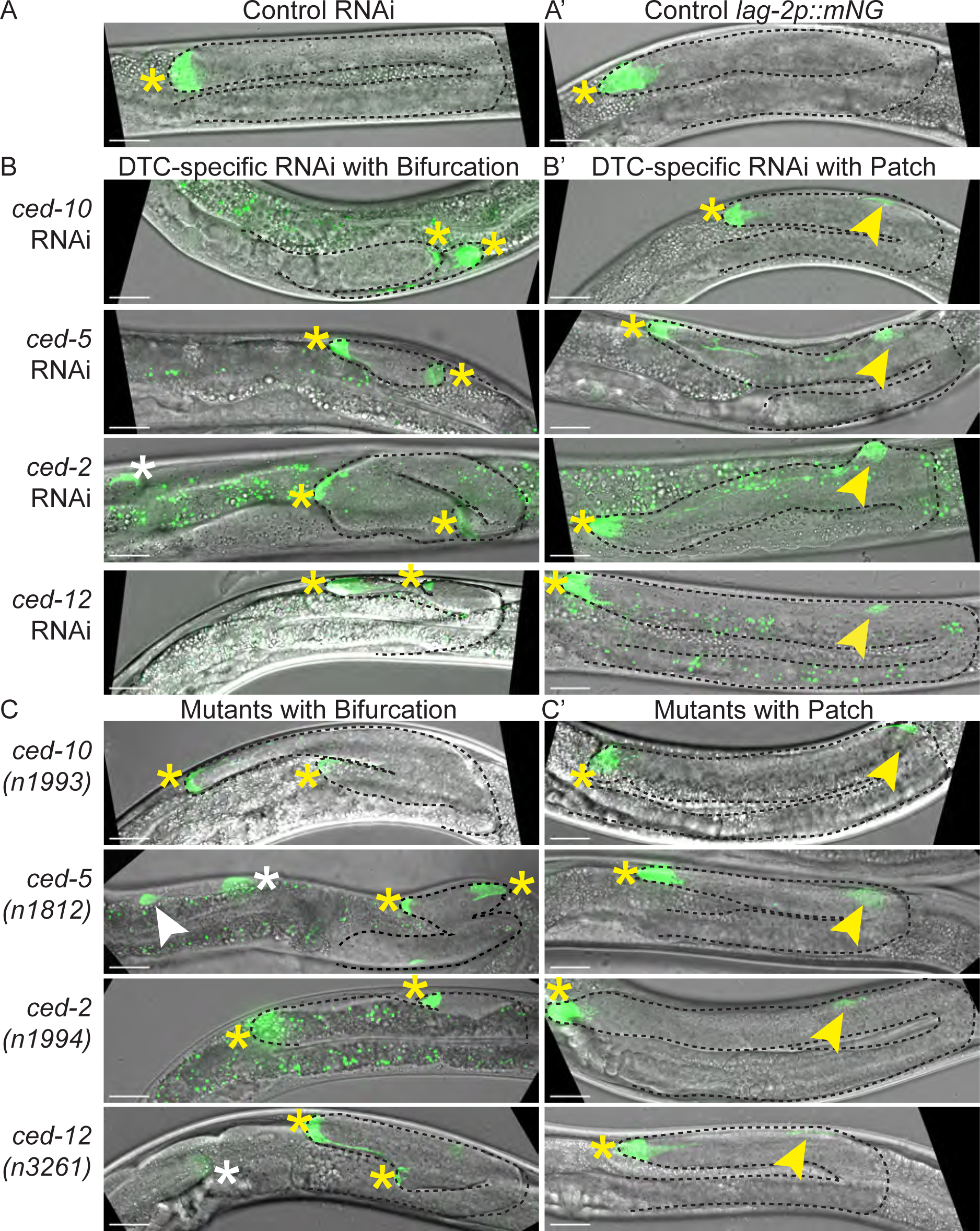
Bifurcation is a DTC cell-autonomous defect caused by Rac-pathway loss of function, and a DTC-membrane marker reveals associated somatic gonad cell morphology defects. **(A)** Strain expressing a combined *lag-2* promoter-driven membrane marker and *rde-1* rescue transgene *lag-2p::mNG::PLC^δPH^::F2A::rde-1* in a genetic background that is *rde-1(ne219)* loss of function and *rrf-3(pk1426)* hypersensitive to RNAi^65^. On control RNAi (empty vector L4440), one site of strong mNG expression is visible in each gonad arm in late L4 hermaphrodites–the distal tip cell. **(A’)** Otherwise wild-type N2 strain bearing *cpIs122[lag-2p::mNeonGreen::PLC^δPH^]*^65^ marker of the DTC. **(B)** DTC-specific RNAi knockdown of Rac-pathway *ced* genes causes a range of cellular and anatomical gonad defects: gonad bifurcation, in which mNG expression is always observed on both tips, and the formation of a second “patch” of mNG expression near the bend of an otherwise anatomically normal gonad **(B’)**. **(C)** Mutants of the Rac-pathway *ced* genes with the *cpIs122* marker of the DTC manifest the same defects as animals treated with DTC-specific RNAi to those genes: gonad bifurcation, in which mNG expression is always observed on both tips, and the formation of a second “patch” of mNG expression near the bend of an otherwise anatomically normal gonad **(C’)**. Maximum projection of GFP fluorescence channel through Z-slices with mNG signal merged with maximum projection of DIC image through slices capturing the gonad tip(s). Imaged at late L4 stage. Visible gonad outlined in black dashed line. Yellow asterisks mark gonad tips of focal gonads, white asterisk marks tip of other gonad arm if visible. Yellow arrowhead marks the patch, and white arrowhead marks patch of other gonad arm if visible. Autofluorescence of the gut is visible as green punctae; this is unrelated to expression of the fluorescent protein. Scale bars 20 μm.

We next examined the DTC in the mutant animals. A cell membrane marker that is specific for the DTC (*cpIs122[lag-2p::mNeonGreen::PLC^δPH^]*^65^ shows a single locus of expression in each gonad arm in otherwise wild-type animals (Figure 2A’). In c*ed-10(n1993), ced-5(n1812), ced-2(n1994),* and *ced-12(n3261)* mutants, this marker reveals the presence of a patch of mNG fluorescence on both tips of bifurcated gonads (Figure 2C) and near the bend in gonads that do not bifurcate (Figure 2C’). The second site of mNG expression appears with high penetrance across mutants (57-80% of mutant animals, N=107), and is always present when we see gonad bifurcation (N=14/14 bifurcated mutant gonads with the DTC marker). Put another way, every bifurcated gonad we observed has a second mNG+ patch, but not every gonad with an ectopic mNG+ patch bifurcates. On the other hand, we did not observe a patch every time we observed DTC pathfinding defects, suggesting these two defects reflect different aspects of Rac pathway function in the DTC. We therefore conclude that Rac-pathway loss-of-function causes the formation of this mNG+ patch, and hypothesize that the patch sometimes causes gonad bifurcation.

### Somatic gonad defects appear during or just after DTC turning

We next sought the origin of the ectopic mNG+ patch. We examined c*ed-10(n1993), ced-5(n1812), ced-2(n1994),* and *ced-12(n3261)* mutants expressing the *cpIs122[lag-2p::mNeonGreen::PLC^δPH^]*^65^ cell membrane marker during larval development. Early in gonad migration (Phase I: ventral migration), a single site of mNG is visible at the growing tip of each gonad arm in wild-type and mutant gonads (Figure 3). As the DTC makes its turns or just after (Phase II: turning), a second mNG+ patch appears in the mutant gonads, attached to the DTC by a filament. Because the patch is connected to the DTC at its first appearance, we hypothesize that the DTC is the source of the patch. Patches are observed in all four mutants by Phase III: dorsal migration.

**Figure 3.**
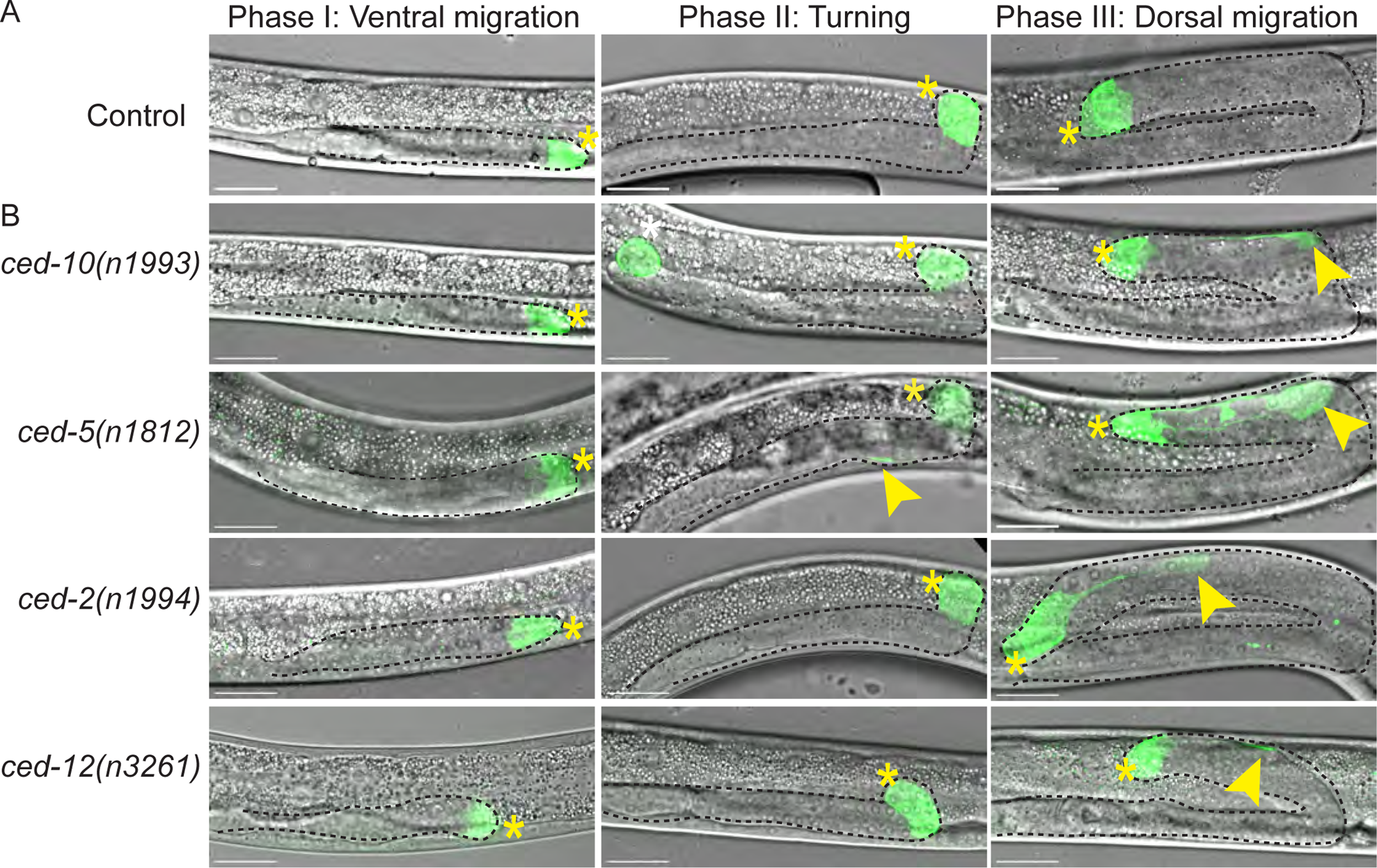
Somatic gonad cell morphology defects manifest at or just after DTC turning. **(A)** Developmental time series of otherwise wild-type N2 animals bearing *cpIs122[lag-2p::mNeonGreen::PLC^δPH^]*^65^ marker of the DTC during ventral DTC migration (left), turning (center), and dorsal migration just after turning (right). The DTC has a gumdrop shape with only very thin trailing filaments. **(B)** Rac-pathway *ced* mutants expressing the same *cpIs122* marker of DTC at the same stages. Patches appear at or after the turning stage. Maximum projection of GFP fluorescence channel through all Z-slices with focal mNG signal merged with maximum projection of DIC image through slices capturing the gonad tip(s). Visible gonad outlined in black dashed line. Yellow asterisks mark gonad tips of focal gonads, white asterisk marks tip of other gonad arm if visible, yellow arrowheads mark the patch. Scale bars 20 μm.

### The DTC fragments after Rac-pathway *ced* loss of function

We tested several hypotheses about how the DTC might give rise to the patch. A bifurcated gonad phenotype has been observed after RNAi treatments that interfere with the cell cycle^63,64^, which cause the normally postmitotic DTC to divide aberrantly, or another somatic gonad cell to be mis-specified as an extra DTC^62,66^. If the second focus of mNG is caused by aberrant cell division or misspecification of another gonad cell, the “patch” would be a second cell with DTC identity. We also considered that overexpression of membrane fluorescent transgene-encoded proteins could cause abnormal cell growth^67^; in that case, strains without such a transgene should not form a patch. Finally, we considered that the DTC breaks apart without dividing, and the patch is a membranous fragment of cellular material.

We first tested whether the patch was or was not a second distal tip cell. We examined a strain expressing an allele encoding endogenously tagged GFP::HLH-2 protein^68^, a tagged transcription factor that localizes to the nucleus of the DTC (among other cells) in control gonads, with some dim cytoplasmic DTC fluorescence signal also observed (Figure 4A). We exposed the strain bearing this allele to *ced-5* and *ced-12* RNAi (as these were the most efficient RNAi treatments for causing gonad bifurcation) and observed GFP::HLH-2 expression in bifurcated gonads. If we detected a bright focus of nuclear signal of this tagged protein in both bifurcated gonad tips, we would conclude that the DTC divided or a second somatic gonad cell became mis-specified as a DTC. However, in 19/19 animals with bifurcated gonads, we observe nuclear localization in only one of the two structures at the tips of the bifurcated gonads and dim cytoplasmic expression in the other (Figure 4B-C). Of note, the tip containing the nucleus of the DTC is not always the tip closest to the correct anatomical position of the DTC (Figure S2). Because GFP::HLH-2 is not encoded by an overexpressed transgene, nor does it encode a membrane-localized fluorescent protein, this experiment also rules out the possibility that driving an overexpressed fluorescent membrane protein in the DTC is responsible for the formation of the structure at the second tip.

**Figure 4.**
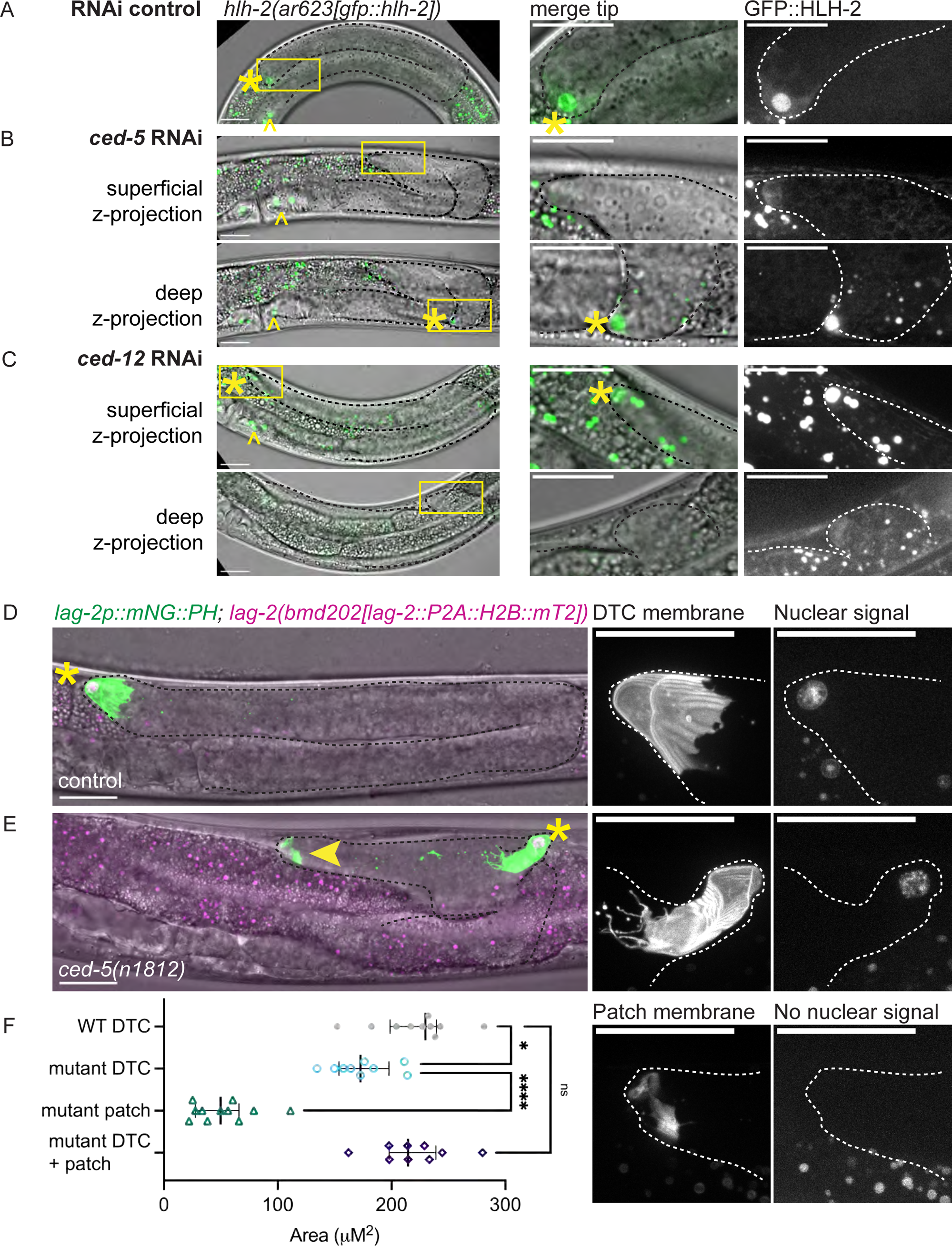
The second tip is not led by a cell, but by a DTC fragment. **(A-C)** A strain expressing an endogenously tagged *hlh-2(ar623[gfp::hlh-1])* allele encoding GFP::HLH-2 transcription factor^68^ on control and experimental whole-body RNAi. Left, fluorescence images merged with DIC projection through gonad tip(s). Center, inset of gonad tip. Right, GFP::HLH-2 alone. Yellow asterisks mark DTC; yellow carats mark proximal gonad cells expressing GFP::HLH-2. **(A)** GFP::HLH-2 strain on control RNAi (empty vector L4440) has concentrated fluorescence in the DTC nucleus and dimmer, diffuse cytoplasmic GFP in the DTC. **(B)** RNAi knockdown of *ced-5* in the strain bearing the *gfp*::*hlh-2* allele. Upper left, maximum projection of GFP fluorescence channel through the more superficial tip of a bifurcated gonad in GFP merged with DIC. Inset right, the more superficial tip has diffuse cytoplasmic GFP::HLH-2 signal. Below, maximum projection of GFP fluorescence channel through the deeper tip of same bifurcated gonad merged with DIC. Inset right, the deeper tip has nuclear GFP::HLH-2 signal. Only animals with a second site of HLH-2::GFP expression were imaged of which N=9 bifurcated gonads. **(C)** RNAi knockdown of *ced-12* in the strain bearing the *gfp*::*hlh-2* allele. Upper left, maximum projection of GFP fluorescence channel through the more superficial tip of a bifurcated gonad in GFP merged with DIC. Inset right, the more superficial tip has nuclear GFP::HLH-2 signal. Below, maximum projection of GFP fluorescence channel through the deeper tip of same bifurcated gonad merged with DIC. Inset right, the deeper tip has diffuse cytoplasmic GFP::HLH-2 signal. N=10 bifurcated gonads. **(D-E)** A strain bearing a transgene that marks the membrane of the DTC, *cpIs122[lag-2p::mNeonGreen::PLC^δPH^],* and a nuclear marker inserted at the endogenous *lag-2* locus (*lag-2::P2A::H2B::mT2)*^69^ in otherwise (D) wildtype and (E) *ced-5(n1812)* genetic backgrounds. Left, maximum projections through all Z-slices with focal mNG (green) and/or mT2 (magenta) signal merged with DIC projection through gonad tip(s). Center, inset of tip, *lag-2p::mNG::PLC^δPH^* only. Right, inset of tip, *H2B::mT2* only. **(D)** Otherwise wild-type strain with DTC membrane and nuclear markers. **(E)** Mutant with bifurcated gonad *ced-5(n1812)* with DTC membrane and nuclear markers. Inset, top, the DTC membrane (center) and nuclear (right) signal. Inset, bottom, the membranous fragment (center) lacks histone signal (right). Visible gonad outlined in black or white dashed line. Yellow asterisks mark gonad tip with DTC nucleus, yellow arrowhead marks the second tip. Yellow boxes show position of insets in larger image. Autofluorescence of the gut is visible as punctae; this is unrelated to expression of the fluorescent proteins. Imaged at L4 stage (L4.6-L4.9), based on vulval morphology^106^. Scale bars 20 μm. **(F)** Graph showing that both the *ced-5(n1812)* DTC and patch are smaller than wild-type DTCs, but the sum of their projected area is not different from that of wild-type DTCs. WT DTC N=10, mutant DTC N=9, mutant patch N=11, mutant DTC + patch N=9. Graphed data is presented with median and interquartile range. One-way ANOVA testing the effect of structure type on size. F_3.00, 32.26_= 70.17, p<0.0001. Dunnett’s T3 multiple comparisons test found that the mean value of structure area was significantly different between WT DTC vs. mutant DTC (p= 0.0126, 95% CI = [9.587, 85.10]) and mutant DTC vs. mutant patch (p < 0.0001, 95% CI=[90.45, 153.6]). However, the structure area was not significantly different between the WT DTC vs. mutant DTC + patch (p=0.9990 95% CI=[-39.54, 43.41]).

We observed the same pattern in a strain coexpressing a *lag-2(bmd202[lag-2::P2A::H2B::mT2])* histone tag^69^ and *lag-2p::mNG* membrane marker (*lag-2p::mNeonGreen::PLC^δPH^*). In an otherwise wild-type genetic background, the DTC membrane and nucleus are the only sites of expression of these markers in the distal gonad (Figure 4D). After crossing these markers into a *ced-5(n1812*) background, nuclear expression is visible in only one of the two membranous bodies (Figure 4E).

We used this same experiment to quantify the sizes of the DTC and enucleate patch appearing in *ced-5(n1812)* mutant animals coexpressing the *lag-2::P2A::H2B::mT2* histone tag and the *lag-2p::mNG* membrane marker vs. that of the otherwise wild-type DTC in L4 animals. After fragmentation, *ced-5* mutant DTCs are smaller in their projected area than control DTCs, with the patches smaller still (Figure 4F). However, the summed sizes of the *ced-5* mutant DTC and patch are not significantly different from the sizes of wild-type L4 DTCs, suggesting that we are seeing two pieces of a fragmented DTC (Figure 4F). We conclude that the ectopic patch is not a daughter of the DTC nor a mis-specified second DTC in the affected gonad arm—it is a cell fragment. This led us to ask how a fragment of cellular material can be competent to lead gonad outgrowth.

### Both fragments of the DTC localize integrin, required for DTC migration, throughout larval development

We next sought to characterize the molecular repertoire of the structures that result from DTC fragmentation. We hypothesized that the cell fragment must localize key regulators of both DTC leader cell and stem cell niche functions in order to lead to the bifurcation of the gonad. Integrin, a receptor that mediates cell interactions with extracellular matrix, is required during DTC migration^4,56,58^. We examined a strain with an endogenously tagged INA-1::mNG allele^70^, which is localized to the surfaces of many cells, including the DTC (Figure 5A). We asked if the patch was decorated by INA-1::mNG protein. We treated the strain with whole-body *ced-5* RNAi to induce DTC fragmentation and imaged gonads with a second focus of INA-1::mNG expression at or near the gonad bend. In the L4 stage, the smaller DTC and the ectopic patch both continue to localize endogenously tagged INA-1::mNG on their surfaces (Figure 5B and 5C). This is true whether the DTC is the fragment that migrates in the correct (Figure 5B) or incorrect (Figure 5C) direction. The ectopic patch therefore contains at least some of the cellular machinery that is known to be required for DTC migration.

**Figure 5.**
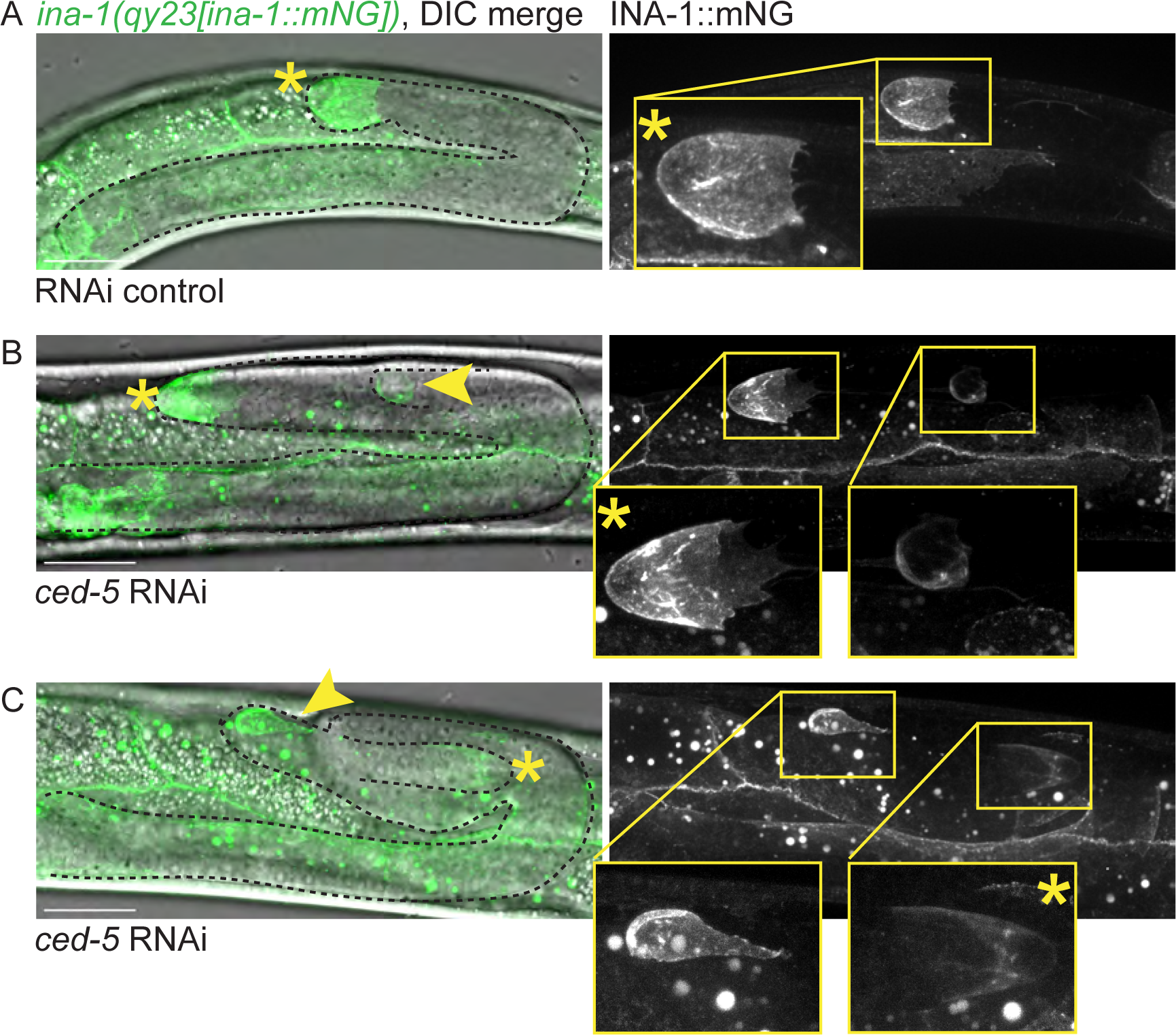
The DTC fragment retains INA-1, a key regulator of gonadogenesis. A strain expressing an endogenously tagged allele of *ina-1(qy23[ina-1::mNG])* has mNG signal on the surface of many cells, including the DTC. Left, merged images of maximum projection through Z-slices with mNG signal and DIC image showing gonad tip(s). Right, GFP channel only. Insets, gonad tip(s). **(A)** INA-1::mNG-expressing strain on control RNAi (empty vector L4440). **(B)** Specimen of INA-1::mNG-expressing strain on *ced-5* RNAi. Note that the DTC is closer to its correct anatomical position than the patch. **(C)** Another specimen of INA-1::mNG-expressing strain on *ced-5* RNAi. Note that the DTC is migrating in the wrong direction with the patch closer to the correct anatomical DTC position; the DTC is deeper in the specimen and is therefore dimmer. Visible gonad outlined in black dashed line. Yellow asterisks mark gonad tip with DTC nucleus, yellow arrowhead marks the enucleate patch. Yellow boxes show positions of insets in larger images. Autofluorescence of the gut is visible as punctae; this is unrelated to expression of the fluorescent protein. Imaged at the L4 stage. Scale bars 20 μm.

### Both DTC fragments localize the stem cell niche stemness signal in larvae but only one maintains expression in adults

LAG-2 is a ligand that activates the GLP-1/Notch receptor on germ cells, acting as the stemness cue, and is necessary for germ cell mitosis throughout gonad migration and adulthood^8,71,72^. Germ cell mitosis provides the pushing force necessary for gonad elongation^4^, so we hypothesized that in order for a gonad to bifurcate, both branches must support germ cell proliferation with a source of LAG-2, at least for a time. We observed L4 animals of a strain expressing endogenously tagged LAG-2::mNG and *lag-2p::myr::TdTomato*^73^ to mark the DTC membrane (Figure 6A, top). We treated this strain with whole-body *ced-5* RNAi to induce DTC fragmentation and investigated LAG-2::mNG localization (Figure 6B). Indeed, LAG-2::mNG signal was apparent on both structures that were positive for the *lag-2p::myr::TdTomato* membrane marker upon RNAi knockdown of *ced-5* (Figure 6C), however the patch has less total LAG-2::mNG than the DTC (Figure 6F). While the “patch” is not a cell with DTC identity, it localizes a protein that can maintain stemness in nearby germ cells.

**Figure 6.**
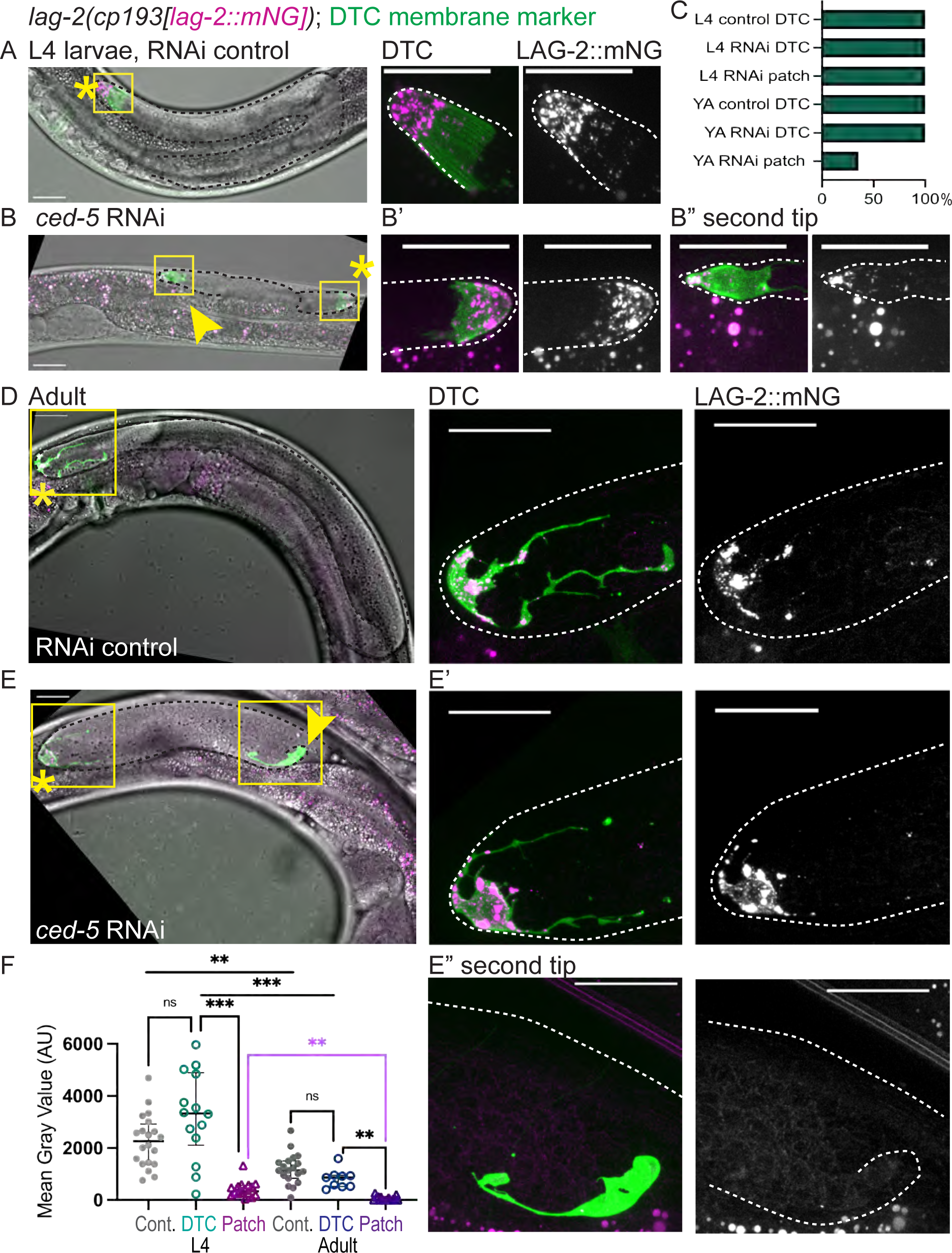
The DTC fragment retains the germline stem cell niche stemness cue in larvae but not adults. **(A)** Top, larval gonad of an animal from a strain expressing an endogenously tagged allele of *lag-2(cp193[lag-2:: mNeonGreen^3xFlag])* as well as a transgene to mark the DTC, *qIs154(lag-2p:: myr::tdTomato)* on control RNAi (empty vector L4440) shows LAG-2::mNG protein (magenta) in the DTC (membrane in green). Left, merged images of maximum projection through Z-slices with mNG and/or TdTomato signal and DIC image showing gonad tip(s). Center, inset of DTC, merged fluorescence channels only. Right, LAG-2::mNG channel alone. **(B)** Larval gonad of an animal from the strain expressing endogenously tagged LAG-2 (magenta) and a DTC membrane marker (green) on *ced-5* RNAi. Note that both structures expressing the DTC membrane marker also have LAG-2::mNG protein on their surfaces. **(B’)** Inset showing merged fluorescence in the DTC (left) and LAG-2::mNG (right). **(B”)** Inset showing merged fluorescence and LAG-2::mNG expression in the second tip. **(C)** Graph showing incidence of LAG-2::mNG signal in the DTC or patch/second tip at L4 stage and in Adults in control and RNAi-treated animals. L4 control DTC, N = 20; L4 RNAi DTC, N = 14; L4 RNAi patch = 16; Adult control DTC, N = 19; Adult RNAi DTC, N = 9; Adult RNAi patch = 17. **(D-E)** The same strain and treatments imaged as Day 1 Adults (staging described in Methods). **(D)** Control RNAi (empty vector L4440). All channels merged. Center, inset of DTC, merged fluorescence channels. Right, LAG-2::mNG channel alone. **(E)** Same strain on *ced-5* RNAi. All channels merged. **(E’)** Center, inset of DTC, merged fluorescence channels. Right, LAG-2::mNG channel. **(E”)** Center, inset of second tip, merged fluorescence channels. Right, LAG-2::mNG channel. Yellow asterisks mark DTC, yellow arrowheads mark the second tip. Yellow boxes show positions of insets in larger images. Autofluorescence of the gut is visible as punctae; this is unrelated to expression of the fluorescent proteins. Scale bars 20 μm. **(F)** Graph showing quantification of LAG-2::mNG fluorescence Mean Gray Value (arbitrary units) in control and *ced-5* RNAi animals in the DTC or patch/second tip. L4 control DTC, N = 20; L4 RNAi-treated DTC = 14; L4 RNAi-treated patch = 16; Adult control DTC, N = 19; Adult RNAi-treated DTC, N= 9; Adult RNAi-treated patch = 17. Graphed data presented with median and interquartile range. One-way ANOVA testing the effect of condition on LAG-2::mNG expression. F_5.000, 28.09_ = 30.14, p <0.0001. Dunnett’s T3 multiple comparisons test found that mean value of LAG-2::mNG expression was significantly different between L4 RNAi DTC vs. L4 RNAi patch (p= 0.0002, 95% CI = [1415, 4243]), L4 RNAi DTC vs. Adult RNAi DTC (p=0.0007, 95% CI = [1002, 3862]), L4 Control DTC vs. Adult Control DTC (p = 0.0024, 95% CI = [307.3, 1823]), Adult RNAi DTC vs. Adult RNAi patch (p = 0.0019, 95% CI = [321.5, 1172]) and highlighted in magenta, L4 RNAi patch vs. Adult RNAi patch (p = 0.0024, 95% CI = [112.3, 586.3]). However, LAG-2::mNG was not significantly different between the L4 Control DTC vs. L4 RNAi DTC (p=0.3631, 95% CI = [-2469, 522.9]) or Adult control DTC vs. Adult RNAi DTC (p=0.2368, 95% CI = [-136.2, 924.0]).

### The ectopic patch does not maintain niche signaling protein long-term, but both DTC fragments persist in adulthood

We interpreted the membrane-localized protein signal on the ectopic patch to come from proteins present at the time of DTC fragmentation or produced soon thereafter. We hypothesized that the enucleate patch has a minimal ability to synthesize new proteins because it lacks a genome. We tested whether the patch can sustain stem cell niche function by examining LAG-2::mNG expression in adult animals. In control young adults at the onset of oogenesis, the LAG-2::mNG protein localizes to the DTC body and its processes (Figure 6D). In young adult *ced-5* RNAi-treated animals, we first observed the *lag-2p::myrTdTomato* membrane-marking signal decorating bifurcated gonad tips and patches perdures (Figure 6E, 6E” and 6C). The membranous cell fragment is not eliminated. However, LAG-2::mNG protein signal is no longer detected in this structure in more than half of adult animals (Figure 6C), and overall LAG-2::mNG expression is significantly decreased in adult patches relative to L4 patches (Figure 6F, magenta bar). The *ced-5* RNAi-treated animals maintain LAG-2::mNG in the DTC itself at levels indistinguishable from control DTCs (Figure 6E’ and 6C), from which we draw two conclusions. First, the loss of LAG-2::mNG signal from the DTC fragment is not due to a global failure of *ced-5* RNAi-treated animals to produce LAG-2::mNG. Second, the DTC that fragmented in the larval stage persists and is both morphologically and molecularly normal hours to days after it breaks apart, suggesting a surprising capacity for cellular recovery in the DTC (Figure 6E’, 6F).

### Bifurcated germlines are mispatterned, with differentiating germ cells at the second tip

Since LAG-2::mNG signal is reduced on the patch relative to the DTC in L4 and is reduced on the surface of the patch in adults (Figure 6F), we predicted that one branch of bifurcated gonads would fail to support stem-like germ cells in later adulthood. We studied germline patterning in branched gonads compared to controls using a germline-specific promoter-driven fluorescent histone *(naSi2(mex-5p::H2B::mCherry))*^74^ to visualize the nuclear morphology of germ cells in otherwise wild-type and *ced-5(n1812)* mutant animals.

A wild-type adult hermaphrodite germline has a distinctive pattern of germ cell nuclear morphology (Figure 7A) extending through the formation of oocytes (Figure 7B). Distal germ cells have small, uniform nuclei (Figure 7C) and visible mitotic figures; these are the cells of the proliferative zone. Proximal to these cells are cells of the transition zone, which have distinctive crescent-shaped nuclear fluorescence (Figure 7D); these are at the leptotene/zygotene stage of meiosis I^14^. Proximal to these cells are meiotic germ cells with “bowl of spaghetti” nuclear morphology of meiotic pachytene^14^ (Figure 7E). As germ cells differentiate into oocytes, the fluorescence signal reveals condensed bivalent chromosomes of diakinesis, and in DIC we observe the cells becoming larger and more cytoplasm-rich (Figure 7B-B”).

**Figure 7.**
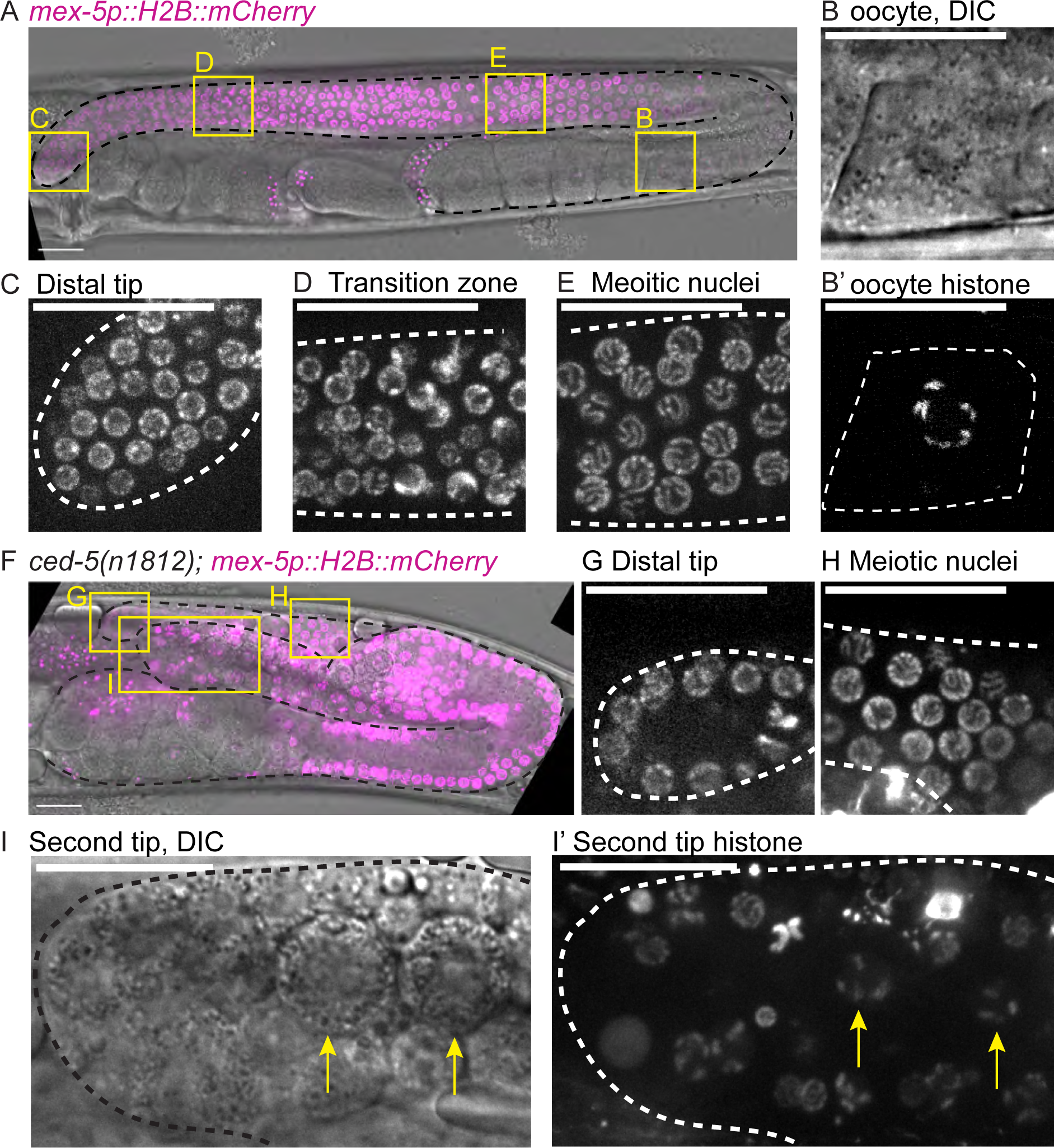
Bifurcated germlines are mispatterned and have differentiating germ cells at the distal end. Micrographs comparing features of adult germlines between control (A-E) and *ced-5(n1812)* (F-I) animals. Germ cell nuclei visualized with the *naSi2(mex-5p::H2B::mCherry)* transgene^74^, magenta in merge. **(A)** Z-projection through DIC image of two-day adult (48 hours post L4) control gonad merged with germ cell histone fluorescence. Boxes show positions of insets that follow. **(B)** Enlargement of cellularizing oocyte showing increased cytoplasm in DIC and, single slice through plane of nucleolus **(B’)** condensed bivalent chromosomes at diakinesis. **(C)** Enlargement of distal tip fluorescence image showing undifferentiated germ cell nuclei. **(D)** Enlargement of the fluorescence image of the meiotic transition zone showing its distinctive crescent-shaped nuclear morphology. **(E)** Enlargement of meiotic pachytene nuclei. **(F)** Z-projection through DIC image of 2-day adult *ced-5(n1812)* gonad merged with germ cell histone fluorescence. Yellow boxes show positions of insets in larger images. **(G)** Enlargement of distal tip fluorescence image showing undifferentiated germ cell nuclei. **(H)** Enlargement of meiotic pachytene nuclei (transition zone is obscured in this sample by the second gonad branch). **(I)** Enlargement of single plane through nucleoli of DIC image of second gonad tip of sample shown in (F). Note large, irregular cells inside. **(I’)** Enlargement of Z-projection through fluorescence image of second gonad tip shown in (I). Note abnormal fluorescent bodies including notable condensed, bivalent-like structures (yellow arrows). Gonads outlined in black or white dashed lines. Autofluorescence of the gut is visible as punctae; this is unrelated to expression of the fluorescent proteins. Scale bars: 20 μm.

In branched gonads of 2-day adults, imaging is complicated by the displacement deep into the animal of one of the two gonad branches. However, in specimens in which both tips are visible, we observe one normal looking branch with the expected distal-to-proximal pattern of undifferentiated-to-differentiated germ cell nuclear morphology (Figure 7F-H). In the second tip, we consistently observed bulbous masses and disorganized H2B::mCherry signal (Figure 7F, I, I’). This includes H2B::mCherry signal consistent with the condensing chromosomes of oogenesis (compare Figure 7I’ to 7B’). A normal distal tip germline stem cell niche is not maintained at the second tip, and germ cells appear to be differentiating in this tip. The persistence of the branch despite the collapse of the ectopic niche suggests that the germ cells do not flow from the branch into the “assembly line” of germline output, but instead differentiate in place.

Aberrant patterning is also visible in younger branched gonads. In control L4 larvae (Figure S3A, S3C-D), most of the distal germline is proliferative, with meiotic transition zone nuclei visible only near the bend. In the proximal germline, chromatin condenses for spermatogenesis^14,75^. In branched gonads of *ced-5(n1812)* L4 larvae undergoing spermatogenesis (Figure S3B), we observe two tubes of germ cells, one of which has the expected patterning with undifferentiated mitotic cells distal (Figure S3E). The second branch tip, however, has germ cells with the crescent nuclear morphology typical of the transition zone of meiotic prophase (Figure S3F). While anatomically it is difficult to identify which is the distal tip and which is the second branch at this stage (see Figure 1), germ cell nuclear morphology reveals that only one branch maintains a stem-like population of germ cells.

As branched gonads lose LAG-2::mNG signal on the patch membrane in adulthood, the germ cells in the second branch become disorganized, lose stemness, and appear to differentiate (taking on crescent-shaped nuclear morphology or showing condensed bivalent chromosomes, and making larger, blocky cells that resemble oocytes). We conclude that, while both DTC fragments resulting from Rac-pathway *ced* loss-of-function are initially competent to lead gonad growth via germ cell proliferation, the enucleate patch cannot sustain adult stem cell niche activity.

## DISCUSSION

During larval development, the *C. elegans* hermaphrodite DTC is both a leader cell and a stem cell niche^1^ and must coordinate cell-biological processes like cell signaling, basement membrane secretion, and adhesion^4^ to form a gonad arm populated by proliferating germ cells. As a leader cell, it is non-protrusive, yet it has long been known that its proper migration requires the canonical “leading edge” Rac pathway members CED-10/Rac1, CED-5/DOCK180, CED-2/CrkII, and CED-12/ELMO. These factors play a role in polarity sensing^40^, however defects in DTC migration polarity have not been reported to cause gonad bifurcation. We found that without Rac pathway function, the DTC fragments during migration into two semi-equivalent structures capable of leading gonad elongation and supporting germ cell proliferation, sometimes driving gonad bifurcation. However, one structure lacks a nucleus and thus lacks the ability to continue to produce LAG-2 protein and thereby act as a stem cell niche through adulthood.

Cell migration depends on major cytoskeletal rearrangements regulated by a complex network of signal transduction pathways involving the Rho-family GTPases, including the highly conserved (in eukaryotes) regulator of lamellipodial cell migration, Rac1^76–78^. Migrating cells often form F-actin-rich protrusions at their leading edge, such as lamellipodia, filopodia, or blebs^38,79^. The DTC curiously does not form protrusions during migration^4,35^. Nevertheless, DTC migration defects have long been observed after loss of function of *ced-10*/Rac1^20–22,29^. Migration events required for branching morphogenesis in the vertebrate lung, kidney, mammary, and salivary glands are also driven by leader cells with a smooth leading edge^80–83^. The mechanisms behind leader cell migration in these processes is largely unknown. Therefore, elucidating regulators of DTC migration could provide a model for vertebrate organogenesis and its misregulation, which occurs in certain cancers^84^.

Rac-pathway factors, including *ced-10*, play many important roles in *C. elegans* development^85^. Rac pathway activity is essential for axon pathfinding^52,86–88^ and dendrite regeneration^89^ of neurons. Rac pathway members have recently been discovered to interact with a CLIC chloride channel protein during the formation of the excretory canal (another tubular organ led by–and in that case composed entirely of–a single cell)^90^. Embryogenesis also requires *ced-10/*Rac in several events, including epidermal cell intercalation^91,92^, gastrulation^93^, mitotic spindle orientation^41^, and of course apoptotic cell corpse clearance^41^. It is not known whether loss-of-function of *ced-10/*Rac or its GEF complex in these other contexts causes cell fragmentation.

Other *C. elegans* cells shed portions of themselves at other stages of development that are cleared by entosis^94^. For example, membranous lobes are either trimmed away or left behind after an entosis event in the primordial germ cells (in a *ced-10*-dependent manner)^95^ and the linker cell (it is not known if this occurs in a Rho-GTPase-dependent manner)^96^. However, these cellular remodeling events are transient, and the residual bodies are eliminated by neighboring cells, whereas the DTC patch is long-lasting and signal-active, at least for a time.

The dividing of a cell without DNA replication into functionally equivalent but genetically distinct daughters (meiosis notwithstanding) is not unprecedented in animals. The superficial epithelial cells of the larval zebrafish skin were recently discovered to divide after terminal differentiation and produce hypoploid descendants^97^. Unlike DTC fragments, however, these descendants each have reduced nuclear content but none lack a nucleus entirely. An example of mouse cells fragmenting has recently been reported in the early embryo as a consequence of ectopic activation of the polar body extrusion pathway in blastomeres^98^. The behavior of the DTC after loss of Rac pathway activity is one of rather few documented instances of cell fragmentation without cell death. How loss of Rac-pathway activity causes the DTC to fragment (and why it only sometimes drives gonad bifurcation) are questions for future study.

The fragmented DTC more closely resembles mutant phenotypes observed in mouse erythrocytes after loss of Rac pathway function. *Rac1^−/−^; Rac2^−/−^* erythrocytes are fragmented and misshapen, and these cells are experimentally determined to be less deformable than cells with Rac1 expression^59^ (the best *C. elegans* BLAST hit of both mouse *Rac1* and *Rac2* is *ced-10*).

Rac1 also functions in human carcinoma cells to downregulate contractile myosin and make cells more deformable; Rac1 dominant-active cells outcompete Rac1-deficient mutant cells via entosis while its depletion causes cells to be stiffer and lose out to more deformable cells^60^. On the other hand, in fibroblasts, Rac1 depletion leads to increased deformability^61^, so Rac1 function has a significant but not strictly directional influence on cell deformability. Taken together, the existing literature and our results suggest that Rac1 is important for cells to dynamically deform without breaking apart. We conclude that, although the DTC does not undergo lamellipodial-driven migration, it nevertheless relies on CED-10/Rac1 to retain its polarity, shape, and integrity at the tip of the growing gonad. Without Rac1 function when cellular cohesion is lost and the cell breaks apart, each fragment can retain leader cell function and support gonad elongation, suggesting that the nucleus is surprisingly dispensable for completion of gonad growth, but is required for long-term maintenance of the stem cell niche.

## METHODS

Sections of this text are adapted from Li et al., 2022^99^, as they describe our standard laboratory practices.

### Strains

Some strains were provided by the CGC, which is funded by NIH Office of Research Infrastructure Programs (P40 OD010440). In strain descriptions, we designate linkage to a promoter with a *p* following the gene name and designate promoter fusions and in-frame fusions with a double semicolon (::). Some integrated strains (xxIs designation) may still contain for example the unc-119 (ed4) mutation and/or the unc-119 rescue transgene in their genetic background, but these are not listed in the strain description for the sake of concision, nor are most transgene 3’ UTR sequences. Complete strain list in Key Resources Table.

### Worm rearing

*C. elegans* strains were kept at 20°C unless otherwise described on standard NGM media and fed *E. coli* OP50. All animals assessed were hermaphrodites, as males have nonmigratory DTCs. Worm populations were synchronized at L1 arrest for developmental staging by standard egg preps^100^.

### Confocal imaging

All images were acquired at room temperature on a Leica DMI8 with an xLIGHT V3 confocal spinning disk head (89 North) with a 63× Plan-Apochromat (1.4 NA) objective and an ORCAFusion GenIII sCMOS camera (Hamamatsu Photonics) controlled by microManager^101^. RFPs were excited with a 555 nm laser, GFPs were excited with a 488 nm laser, and CFP was excited by a 445 nm laser. Z-stacks through the gonad were acquired with a step-size of 1µm unless otherwise noted. Worms were mounted on agar pads with 0.01 M sodium azide.

### RNAi

*E. coli* HT115(DE3) containing the L4440 plasmid with or without a dsRNA trigger insert from the Ahringer^102^ or Vidal^103^ RNAi libraries was grown as a single-colony overnight culture at 37°C, expression induced with IPTG (Apex BioResearch Products, cat# 20-109) for one hour at 37°C, and was plated and allowed to grow at least overnight on NGM plates. Note, Ahringer *ced-10* clone number C09G12.8A has <30 nt homology to the *ced-10* 3’ UTR and yielded inconsistent knockdown; Vidal *ced-10* clone AAA28140 was used preferentially. Worm populations were synchronized by bleaching according to a standard egg prep protocol^100^, plated on NGM plates seeded with RNAi-expressing bacteria as arrested L1 larvae, and kept on RNAi until the time of imaging. Whole-body RNAi treatment was conducted at 20°C, and DTC-specific RNAi treatment was conducted at 16°C due to a temperature-sensitive *rrf-3* mutation in the DTC-specific RNAi strain. Whole-body *ced-12* RNAi on the *hlh-2(ar623[gfp::hlh-2])* strain included specimens imaged after two generations on RNAi.

DTC-specific RNAi was performed using strain NK2115^65^, a genetic background carrying an *rde-1(ne219)* loss of function that prevents RNAi activity globally, with RNAi function restored in the DTC (in an operon along with a coding sequence for membrane-tethered mNeonGreen) by a transgene *lag-2p::mNG::PLC^δPH^::F2A::rde-1* and a *rrf-3(pk1426)* mutation that enhances RNAi.

### Image analysis

Images were processed in FIJI^104^ (Version: 2.14.1/1.54f), and images that required stitching used the ImageJ Stitching Plugin^105^. Detailed descriptions of image analysis for different experiments are provided below.

### Measurement of DTC vs patch area

To compare the area of the WT DTC with the mutant DTC and enucleate patch, WT and mutant gonads coexpressing the *lag-2p::mNeonGreen::PLC^δPH^* membrane marker and *lag-2::P2A::H2B::mT2* nuclear marker were imaged with a 1.0 µm Z-step size. Only late L4 animals were imaged for both WT and mutants and only mutants with a patch or bifurcation (both marked by a second focus of mNG expression) were imaged. Late L4 animals were defined as between the L4.6-L4.9 larval stage, based on vulval morphology^106^. Using FIJI^104^, maximum intensity z-projections were made through the depth of the entire DTC and patch structures. The structures were outlined by hand (based on membranous mNG signal in the GFP channel) using the freehand selections tool to define ROIs that corresponded to the boundary of the structure. The area of these ROIs was then measured in µm^2^.

### Fluorescence intensity of LAG-2::mNeonGreen

For quantitative comparisons of fluorescence intensity shown in FIGURES, gonads were imaged with uniform laser power and exposure times with 0.3 µm Z-steps. For the L4 stage, only L4.6-L4.9 (late L4) animals were imaged, staged based on vulval morphology^106^. Young Adults were identified as ∼24h post the L4 stage and by the presence of some fertilized embryos. Using FIJI^104^, sum-intensity z-projections were made through the depth of the focal DTC and patch structures as visualized with the TdTomato signal in the RFP channel, excluding deeper Z-slices with appearance of large, bright gut granules in the footprint of the DTC. The structures were outlined with the Threshold function to make ROIs, and Mean Gray Value (and Raw Integrated Density) were measured inside the ROI in arbitrary units. The same ROIs were shifted onto worm gonad background and GFP fluorescence intensity was measured again. This background was subtracted from the signal for the final value of LAG-2::mNG detected in the structure. In the *ced-5* RNAi group, only RNAi-treated specimens with patches or bifurcations were analyzed; escapers were discarded. Because different regions of the gonads in adults are often in different focal planes, not every sample had both a visible DTC and a visible patch. Along those lines, we measure lower LAG-2::mNG signal in adult control and *ced-5* RNAi DTCs compared to L4 DTCs. We attribute this to imaging adult DTCs at a greater tissue depth, which reduces signal capture, and with greater background autofluorescence in adults relative to L4s. Patches in the adult, on the other hand, were usually on the surface of the gonad and were never under the gut. We never observed adult DTCs (either control or *ced-5* RNAi-treated) that lacked LAG-2::mNG signal, while the patches of *ced-5* RNAi-treated adults lacked noticeable signal in more than half of all samples.

### Statistical analyses

Sample sizes, tests, test statistics, and p values are given for each analysis in the accompanying figure legends. All statistical testing was carried out using Prism (GraphPad Prism Version 10.10 (264) for macOS), GraphPad Software, San Diego, CA. Significance bars in each graph were produced by Dunnett’s T3 Test for multiple comparisons of selected groups.

## ACKNOWLEDGEMENTS

We thank Taylor Medwig-Kinney, Pu Zhang, Rob Dowen and his lab members, Brian Kinney, Fred Koitz, and other Gordon Lab members for feedback on the manuscript and sharing resources. Funded by NIGMS Grant 1R35GM147704 to KLG.

---

**Figure S1.**
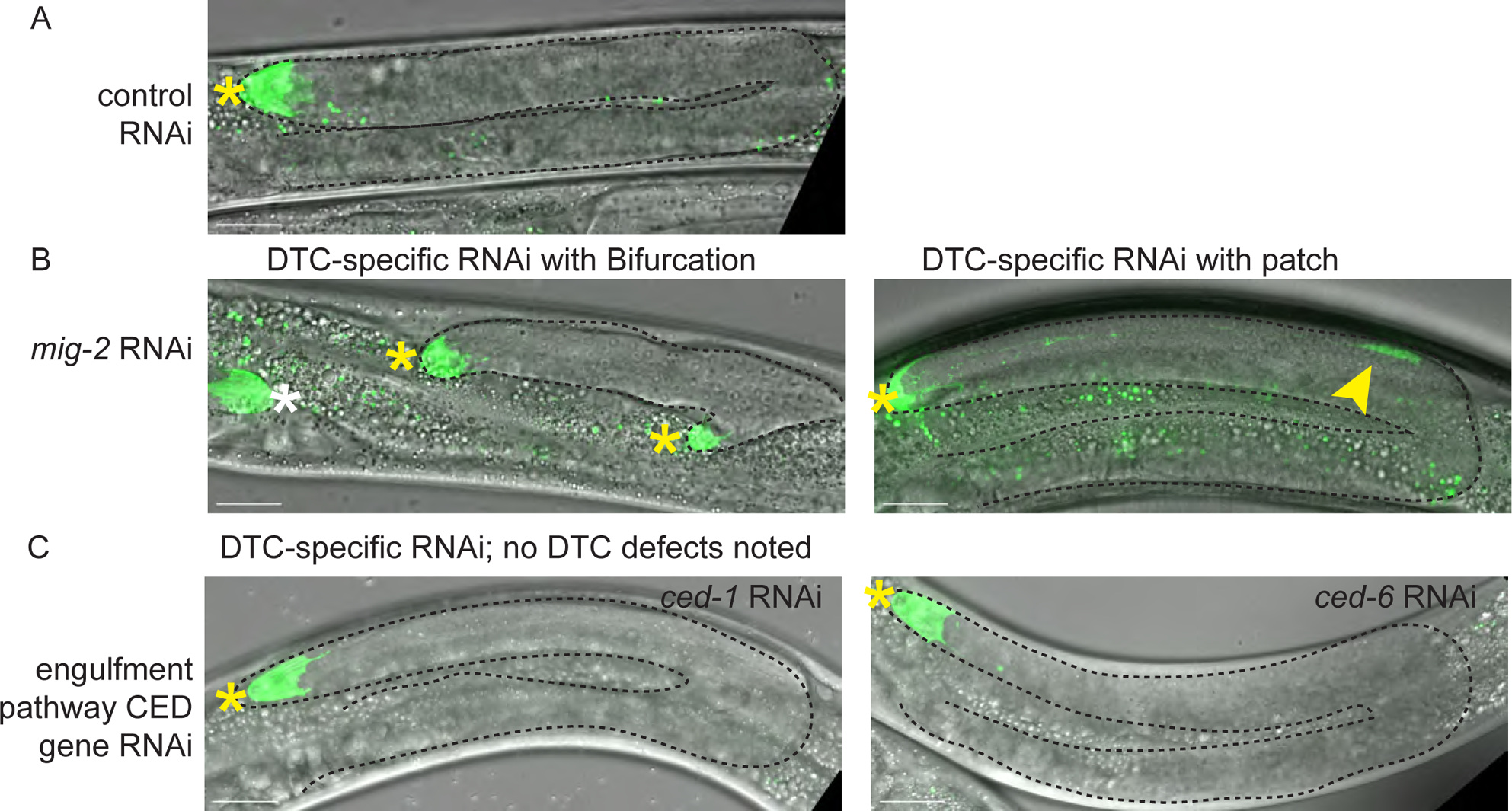
DTC-specific knockdown of *mig-2/*RhoG GTPase, but not parallel cell-engulfment pathway *ced* genes causes DTC cell morphology and migration/bifurcation defects. Strain expressing a combined *lag-2* promoter-driven membrane marker and *rde-1* rescue transgene *lag-2p::mNG::PLC^δPH^::F2A::rde-1* in a genetic background that is *rde-1(ne219)* loss of function and *rrf-3(pk1426)* hypersensitive to RNAi. **(A)** On control RNAi (empty vector L4440), one site of strong mNG expression is visible in each gonad arm in late L4 hermaphrodites–the distal tip cell. No expression is observed near the bend of the gonad (inset). **(B)** DTC-specific RNAi knockdown of the *mig-2/*RhoG gene causes a range of cellular and anatomical gonad defects: gonad bifurcation (left), in which mNG expression is always observed on both tips, and the formation of a second “patch” of mNG expression near the bend of an otherwise anatomically normal gonad (right). N=34 total RNAi-treated animals scored, 7/34 had second patch only and an additional 2/34 had a bifurcated gonad. *mig-2* acts upstream of the Rac pathway *ced* genes^40^. **(C)** DTC-specific knockdown of *ced-1* (N=33) and *ced-6* (N=29), members of a parallel engulfment pathway to *ced-10, ced-5, ced-2,* and *ced-12.* No DTC or gonad morphology defects were observed, though in 3/62 samples a delay in dorsal elongation was noted in a single gonad arm. Maximum projection of GFP fluorescence channel through all Z-slices with mNG signal merged with maximum projection of DIC image through slices capturing the gonad tip(s). Imaged at late L4 stage. Visible gonad outlined in black dashed line. Yellow asterisks mark gonad tips of focal gonads, white asterisk marks tip of other gonad arm, yellow arrowhead marks the patch. Autofluorescence of the gut is visible as green punctae; this is unrelated to expression of the fluorescent protein. Scale bars 20 μm.

**Figure S2.**
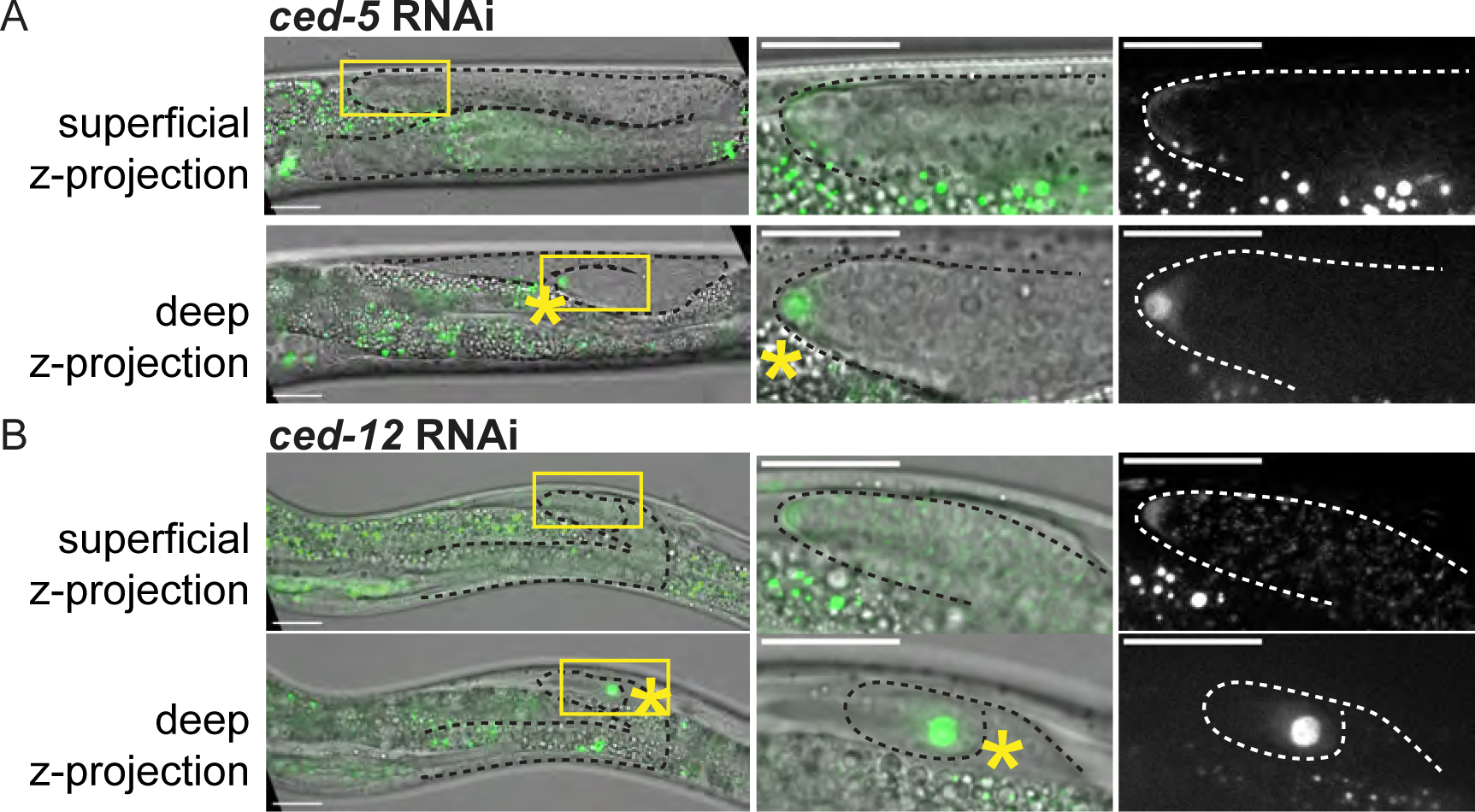
The nuclear DTC signal is not always found in the distal-most tip. **(A)** RNAi knockdown of *ced-5* in the strain bearing the *gfp*::*hlh-2* allele. Upper left, maximum projection of GFP fluorescence channel through the more superficial tip of a bifurcated gonad in GFP merged with DIC. Inset right, the more superficial tip has diffuse cytoplasmic GFP::HLH-2 signal, despite falling in the anatomically correct position of a gonad distal tip. Below, maximum projection of GFP fluorescence channel through the deeper tip of same bifurcated gonad merged with DIC. Inset right, the deeper tip has nuclear GFP::HLH-2 signal. N = 1/9 total bifurcated gonads imaged **(B)** RNAi knockdown of *ced-12* in the strain bearing the *gfp*::*hlh-2* allele. Upper left, maximum projection of GFP fluorescence channel through the more superficial tip of a bifurcated gonad in GFP merged with DIC. Inset right, the more superficial tip has diffuse cytoplasmic GFP::HLH-2 signal. Below, maximum projection of GFP fluorescence channel through the deeper tip of same bifurcated gonad merged with DIC. Inset right, the deeper tip has nuclear GFP::HLH-2 signal, despite having made an extra turn towards the viewer. N = 1/10 total bifurcated gonads imaged. Visible gonad outlined in black or white dashed line. Yellow asterisks mark gonad tip with DTC nucleus, yellow arrowhead marks the enucleate patch. Yellow boxes show position of insets in larger image. Autofluorescence of the gut is visible as punctae; this is unrelated to expression of the fluorescent proteins. Imaged at L4 stage. Scale bars 20 μm.

**Figure S3.**
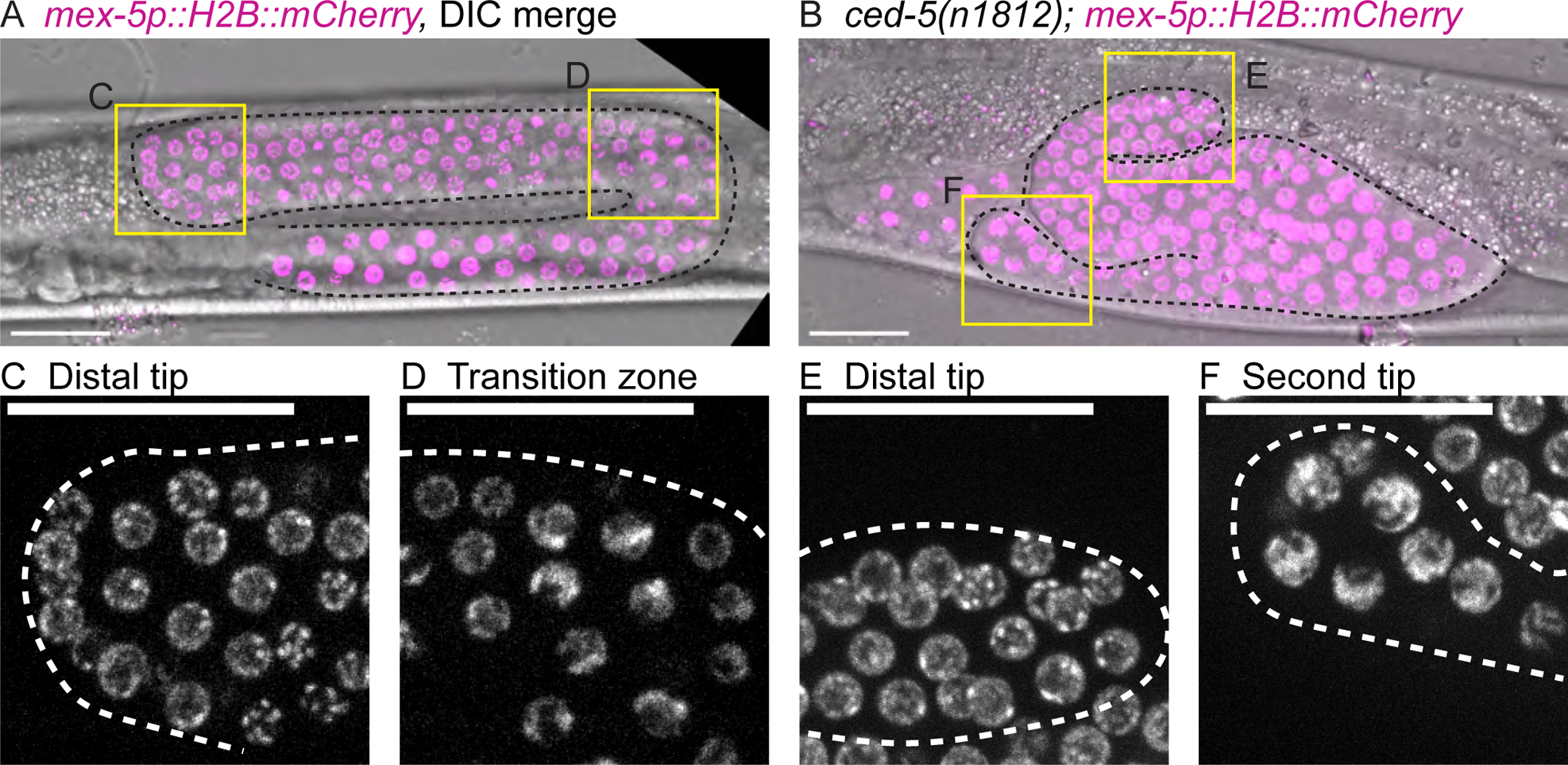
Bifurcated germlines are mispatterned and have differentiating germ cells at the distal end in the L4 stage. Micrographs comparing features of adult germlines between control (A, C, D) and *ced-5(n1812)* (B, E, F) animals. Germ cell nuclei visualized with the *naSi2(mex-5p::H2B::mCherry)* transgene^74^, magenta in merge. **(A)** Z-projection through DIC image of L4 control gonad merged with germ cell histone fluorescence. Boxes show positions of insets that follow. **(B)** Z-projection through DIC image of L4 *ced-5(n1812)* gonad merged with germ cell histone fluorescence. Boxes show positions of insets that follow. **(C)** Enlargement of control distal tip fluorescence image showing undifferentiated germ cell nuclei. **(D)** Enlargement of the fluorescence image of the control meiotic transition zone showing its distinctive crescent-shaped nuclear morphology. **(E)** Enlargement of distal tip of *ced-5(n1812)* fluorescence image of gonad distal tip showing undifferentiated germ cell nuclei. **(F)** Enlargement of second tip of *ced-5(n1812)* mutant goand fluorescence image showing crescent-shaped germ cell nuclei characteristic of cells that have entered meiosis. Gonads outlined in black or white dashed lines. Autofluorescence of the gut is visible as punctae; this is unrelated to expression of the fluorescent proteins. Scale bars: 20 μm.

